# Clinical grade ACE2 effectively inhibits SARS-CoV-2 Omicron infections

**DOI:** 10.1101/2021.12.25.474113

**Authors:** Vanessa Monteil, Devignot Stephanie, Jonas Klingström, Charlotte Thålin, Max J. Kellner, Wanda Christ, Sebastian Havervall, Stefan Mereiter, Sylvia Knapp, Nuria Montserrat, Benedict Braunsfeld, Ivona Kozieradzki, Omar Hasan Ali, Astrid Hagelkruys, Johannes Stadlmann, Chris Oostenbrink, Gerald Wirnsberger, Josef M. Penninger, Ali Mirazimi

## Abstract

The recent emergence of the SARS-CoV-2 variant Omicron has caused considerable concern due to reduced vaccine efficacy and escape from neutralizing antibody therapeutics. Omicron is spreading rapidly around the globe and is suspected to account for most new COVID-19 cases in several countries, though the severity of Omicron-mediated disease is still under debate. It is therefore paramount to identify therapeutic strategies that inhibit the Omicron SARS-CoV-2 variant. Here we report using 3D structural modelling that Spike of Omicron can still associate with human ACE2. Sera collected after the second mRNA-vaccination did not exhibit a protective effect against Omicron while strongly neutralizing infection of VeroE6 cells with the reference Wuhan strain, confirming recent data by other groups on limited vaccine and convalescent sera neutralization efficacy against Omicron. Importantly, clinical grade recombinant human soluble ACE2, a drug candidate currently in clinical development, potently neutralized Omicron infection of VeroE6 cells with markedly enhanced potency when compared to reference SARS-CoV-2 isolates. These data show that SARS-CoV-2 variant Omicron can be readily inhibited by soluble ACE2, providing proof of principle of a viable and effective therapeutic approach against Omicron infections.

## Introduction

The initial step of SARS-CoV-2 infection is binding of the viral Spike protein to Angiotensin converting enzyme 2 (ACE2) [1-3], followed by proteolytic processing of the trimeric Spike protein [4, 5]. Blocking the Spike/ACE2 interaction is the fundamental principle for the activity of neutralizing antibodies induced by all current vaccines [6, 7]. Similarly, all approved and in development antibodies or nanobodies act by blocking the interaction of the cell-entry receptor ACE2 and the viral Spike protein [8]. Interfering with binding of Spike to its surface receptor ACE2 has become a key paradigm of both vaccine design and multiple therapeutic approaches including ACE2 based therapeutics [9-12]. Vaccines and antibody therapeutics have had an enormous impact on the COVID-19 pandemic. However, many variants of SARS-CoV-2 have emerged throughout the pandemic [13, 14], some of which have been designated variants of concern (VOC) by the WHO because of their increased infectivity and transmissibility.

Mutations in the viral Spike protein are of critical importance in viral evolution. These mutations do not only affect the infectivity and transmissibility of SARS-CoV-2, but also reduce the potency of vaccines, convalescent sera, and monoclonal antibody therapeutics [14-20]. The recent emergence of the Omicron variant, which contains 61 nonsynonymous mutations relative to the original Wuhan strain, is a key example [21-23] More such variants will probably evolve in the future, also in part due to population scale measurements and thereby mounting evolutionary pressure on the virus strains. Mathematical modelling to simulate the dynamics of wild-type and variant strains of SARS-CoV-2 in the context of vaccine rollout and nonpharmaceutical interventions has shown variants with enhanced transmissibility such as Delta and Omicron frequently increase epidemic severity, whereas those with partial immune escape either fail to spread widely or primarily cause reinfections and breakthrough infections [24]. However, when these phenotypes are combined, a variant can continue spreading even as immunity builds up in the population, limiting the impact of vaccination and exacerbating the epidemic. Moreover, based on the experience with HIV therapeutics, it is possible that SARS-CoV-2 variants will emerge that reduce the efficacy of RNA polymerase and protease inhibitors[25-27]. It is therefore paramount to identify robust and universal therapeutics for the prevention and treatment of Omicron and future variants of concern.

The Spike/ACE2 interaction is the crucial first step of viral infection. There are concerns that Omicron might also carry mutations that alter its dependency on ACE2 as entry gate, thereby also changing its infectivity and tissue tropism. Here we report, using molecular 3D modelling, that human ACE2 can associate with the receptor binding domain (RBD) and full-length Spike proteins of the Omicron SARS-CoV-2 variant. Sera from SARS-CoV-2 naïve and doubly mRNA vaccinated people fail to neutralize Omicron infections of VeroE6 cells, in line with other emerging data that Omicron can in part escape humoral immunity induced by vaccination and in convalescent people [28-30], explaining the increasing number of breakthrough infections. Most importantly, soluble ACE2/APN01, already being tested in clinical trials for severe (WHO stage 4-6) COVID-19 (NCT04335136) and in a phase 1 inhalation trial for early intervention, potently and effectively neutralizes infections of Omicron. In addition to our recent data that ACE2 blocks all other known SARS-CoV-2 variants of concern, these results provide the blueprint for a universal anti-COVID-19 agent with the potential to alleviate or prevent infections with Omicron.

## Results

### 3D modelling of Omicron Spike and Omicron binding to human ACE2

Many single or compound mutations, especially in the RBD domain of the viral Spike, have been described and either hypothesized or demonstrated to affect binding to the cell entry receptor ACE2 (see [14] for a review). For the newly emerged Omicron variant, it has been proposed, based on clinical presentations and preprint data, that the mutant Omicron Spike might not or only in an altered manner bind to ACE2, a critical issue for the understanding of disease pathogenesis and viral tropism and therefore potential treatment and vaccine designs. Omicron carries 36 mutations in Spike including multiple alterations in the RBD. We first rendered all mutations of the prefusion state Spike of Omicron in 3D using molecular modelling (Figure 1a). The RBD changes of Omicron and the location(s) of the respective mutations are further depicted in a 3D model of the viral Spike RBD domain (Figure 1b).

**Figure 1.**
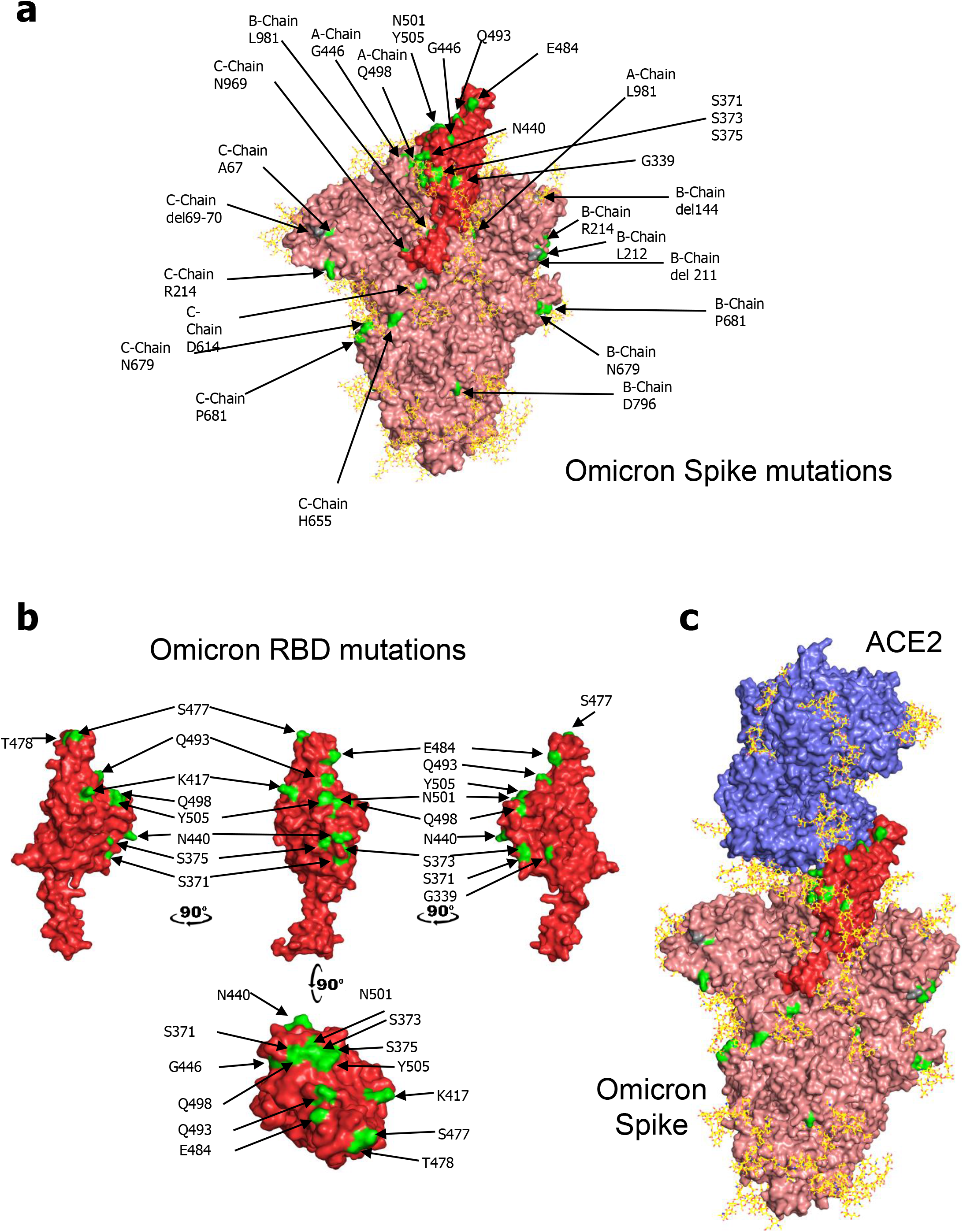
Omicron Spike and RBD mutations and ACE2 interaction. **(a)** PyMOL rendering of the trimeric full-length SARS-CoV-2 Spike protein of the Wuhan reference strain. One RBD domain is shown in red. Indicated in green are positions mutated in the Omicron SARS-CoV-2 variant Spike used in experiments in this study. Depicted in yellow are the glycan-modifications of the Spike protein. **(b)** PyMOL rendering depicting the SARS-CoV-2 RBD domain from the Wuhan reference strain with positions mutated in Omicron RBD highlighted in green. **(c)** PyMOL rendering of trimeric Spike (RBD in red) with human ACE2 (magenta). Positions mutated in Omicron are highlighted in green to indicate their position relative to the interaction surface with ACE2. Glycan modifications are depicted in yellow on both Spike and ACE2 protein.

Importantly, structural modelling of ACE2 binding revealed that pre-fusion Omicron Spike could still associate with human ACE2 (Figure 1c). Mutations at residues K417N, E484A, Q493R, Q498R, N501Y, and Y505H at the RBD directly affect binding of Omicron Spike to human ACE2, resulting most likely in greater binding affinity [31]. Of note, the real affinity of the Omicron Spike-ACE2 interaction needs to be determined in direct biochemical experiments and cannot be deduced faithfully from our modelling. Interestingly, Omicron carries mutations at Q498 at Q493, both of these mutations have been reported in mouse-adapted virus strains [32-34],including our recently developed mouse adapted maVie16 SARS-VoV-2 strain that causes severe C OVID-19 in mice [35]. Thus, it is likely that Omicron will infect rodents. We also modelled the 22 N-glycosylation sites of Spike we and others have previously reported, some of which (N165, N234, N343) directly interact with ACE2 or its glycans[36, 37]. Intriguingly, despite the unprecedented number of observed mutations, none of the N-glycosylation sites critical for ACE2 binding are altered in Omicron Spike (Figure 1c). These molecular modelling data support that pre-fusion Omicron Spike can still readily associate with human ACE2.

### Impaired neutralization of Omicron by RNA vaccine elicited antibodies

We finally tested whether sera from vaccinated people could also still neutralize Omicron infections of VeroE6 cells. To this end, we obtained sera from four SARS-CoV-2 naïve healthcare workers who received an mRNA vaccine (Comirnaty); these sera were collected following ethic approvals 5-7 weeks after the second vaccination. In all cases we observed significant inhibition of infection of the reference SARS-CoV-2 strain. However, at the dilutions used we did not detect any neutralization of Omicron infection of VeroE6 cells (Figure 2). These results are in line with recently emerging studies [38-41] that Omicron carries mutations that can in part escape neutralization by peak antibody levels elicited by the currently standard mRNA vaccines.

**Figure 2.**
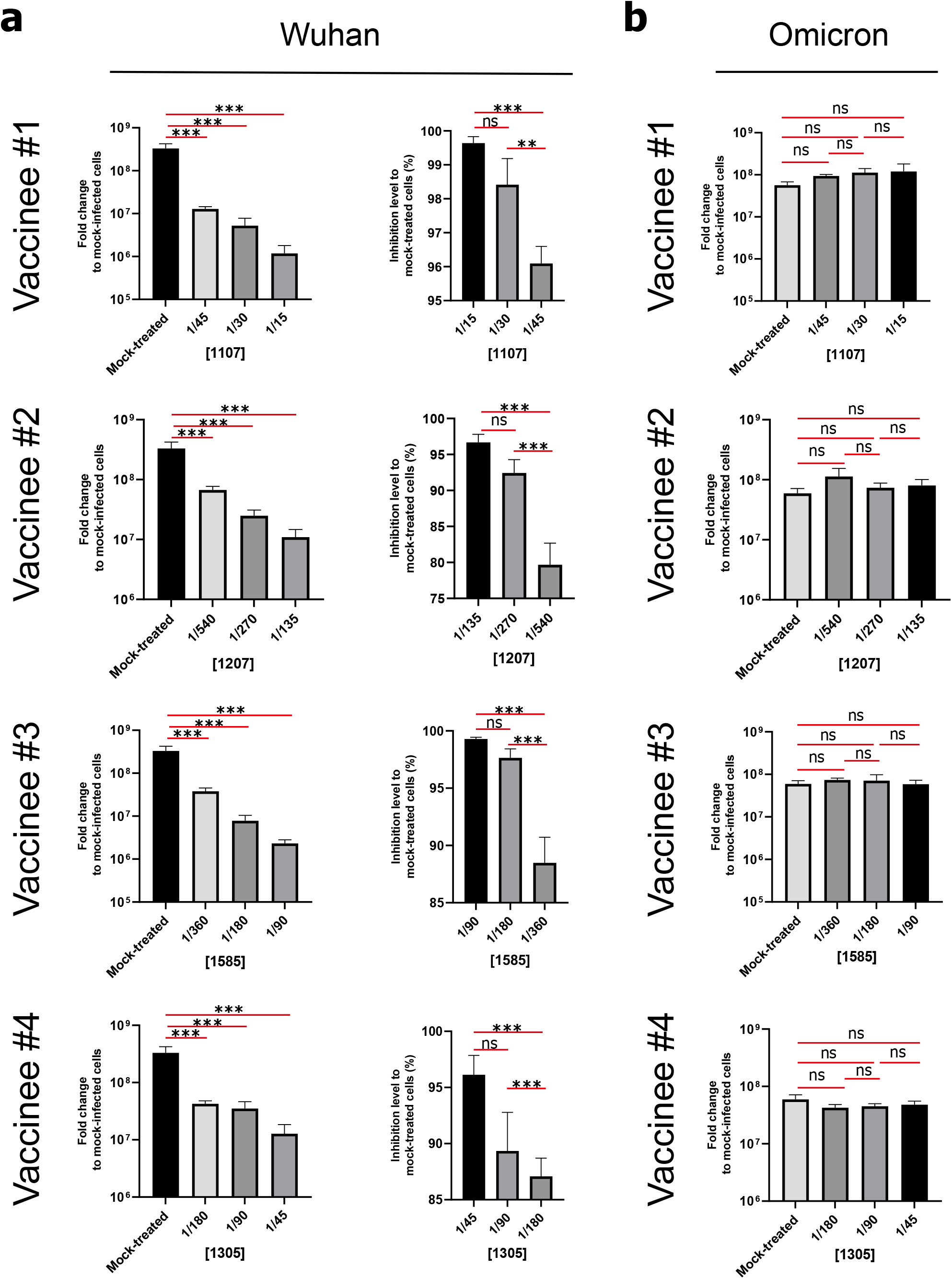
Loss of viral neutralization potency of doubly mRNA vaccinated subjects’ sera against Omicron. **(a)** Diagrams depict the level of infection (left panels) or inhibition of infection (right panels) with the SARS-CoV-2 Wuhan reference strain isolate in the presence of the indicated dilutions of four different mRNA vaccinee’s sera. Sera were taken 5-7 weeks after the second mRNA vaccination, i.e. at the peak of the antibody response, **(b)** Diagrams depict the level of infection with the SARS-CoV-2 Omicron strain isolate in the presence of the indicated dilutions of sera from (a). Data from Vero E6 cell infections are shown at MOI 0.01. Shown in **(a)** and **(b)** are means of triplicate analyses with standard deviations. Statistical significance is indicated by asterisks (p-value < 0.01: **; p-value < 0.001: *** calculated using Two-way ANOVA).

### ACE2/APN01 effectively neutralizes the Omicron SARS-CoV-2 variant

We have previously reported that clinical grade soluble recombinant human ACE2 (APN01) can effectively reduce the SARS-CoV-2 viral load in VeroE6 cells in a dose dependent manner, using a reference virus isolated early during the pandemic[42]. This virus carried the same Spike sequence as the originally reported virus. Moreover, we and others have shown that ACE2/APN01 not only binds significantly stronger to RBD or full-length Spike proteins of all tested variants (alpha, beta, gamma, delta), but also more potently inhibits viral infection by these strains [30, 31].

To test whether APN01 can also neutralize Omicron SARS-CoV-2 isolates, we performed neutralization assays in VeroE6 cells and compared its inhibitory potency side by side to our reference strain. Of note, VeroE6 cells are commonly used to assay SARS-CoV-2 infectivity and drug efficacy. The reference virus was previously reported [42, 43] and carries the Spike amino acid sequence described for the first Wuhan virus isolate. As reported before [42], APN01 markedly reduced viral replication of the SARS-CoV-2 reference strain in a dose dependent manner (Figure 3a). Importantly, the inhibitory potency of APN01 was significantly increased towards the Omicron variant of concern (Figure 3b). These results show that clinical grade soluble human ACE2/APN01 potently blocks SARS-CoV-2 infections of the recently emerged Omicron VOC.

**Figure 3.**
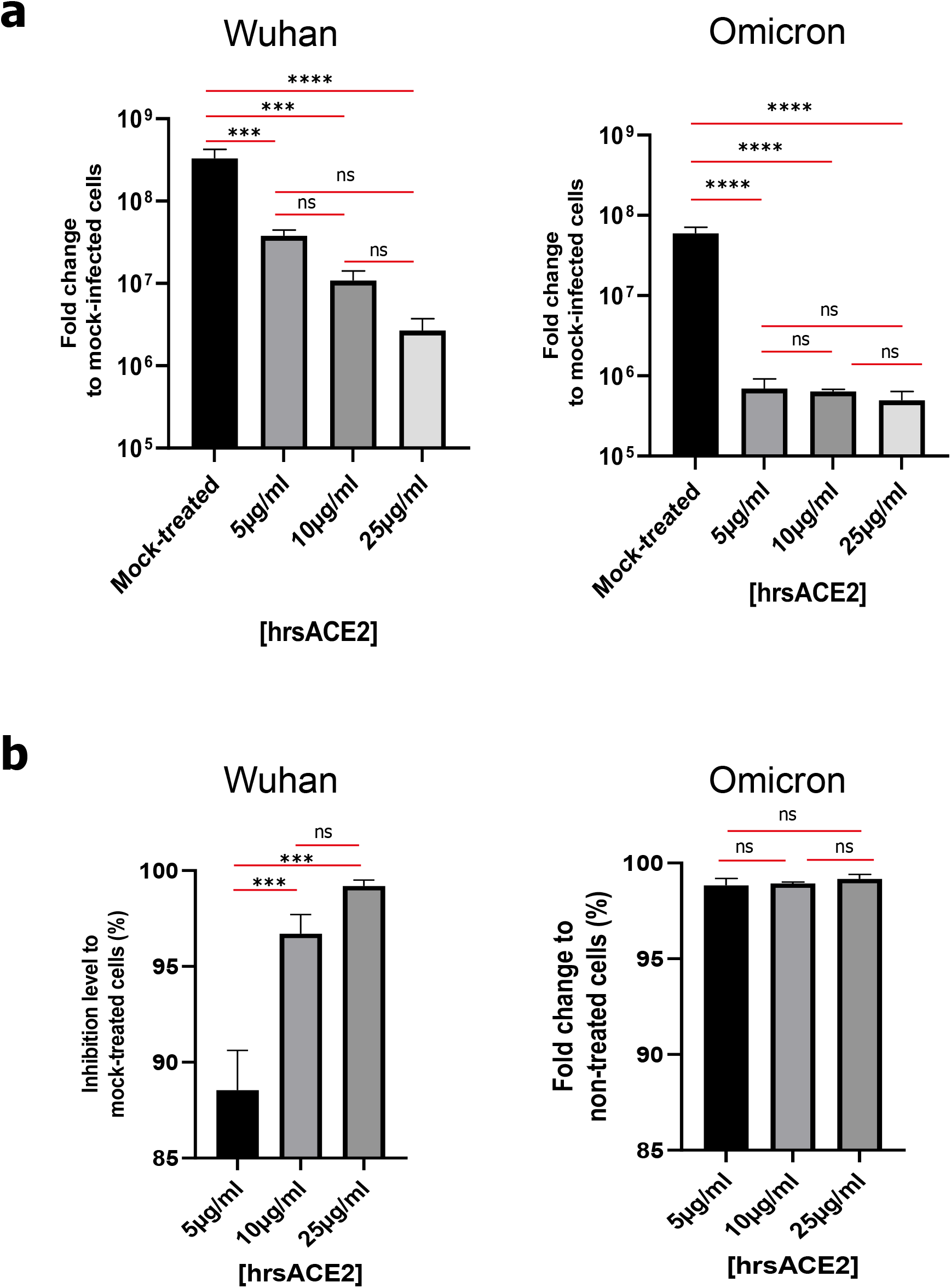
Increased potency of soluble ACE2/APN01 against the Omicron SARS-CoV-2 Variant of Concern. **(a**,**b)** Diagrams depict the level of infection **(a)** and level of inhibition of infection **(b)** of Vero E6 cells with the Wuhan SARS-CoV-2 reference isolate (left panels) and the Omicron SARS-CoV-2 isolate (right panels) in the presence of the indicated concentrations of soluble ACE2/APN01 as compared to mock treatment. Vero E6 cells were infected at MOI 0.01. Shown are means of triplicate analyses with standard deviations. Statistical significance is indicated by asterisks (p-value < 0.001: ***; p-value < 0.0001: **** calculated using Two-way ANOVA).

## Discussion

Seventeen years after the epidemic of SARS coronavirus, the novel coronavirus SARS-CoV-2 emerged, resulting in an unprecedented pandemic. Throughout the COVID-19 pandemic, a plethora of genetic SARS-CoV-2 variants have emerged with some strains displaying increased infectivity and transmissibility, therefore designated Variants of Concern (VOC; https://www.cdc.gov/coronavirus/2019-ncov/variants/variant-info.html and https://www.who.int/en/activities/tracking-SARS-CoV-2-variants/ for further information). Besides the VOC B.1.1.7 (Alpha), B.1.351 (Beta), P.1 (Gamma) and B.1.617.2 (Delta), multiple Variants of Interest (VOI) are also circulating including B.1.526 (Iota), B.1.427, B.1.429, B.1.617.1 (Kappa), B.1.617.3, or B.1.525 (Eta). The last weeks have seen the emergence of the Omicron VOC with an unprecedented number of genetic alterations, including 36 changes in Spike [44] https://www.gisaid.org/hcov19-variants/. Omicron rapidly spreads and readily re-infects doubly and even triply vaccinated people as well as patients recovered from previous SARS-CoV-2 infections, sometimes leading to severe breakthrough COVID-19, as had been previously observed with the Delta variant [21, 22]. Omicron already as a devastating impact on case numbers in several countries, causing lockdowns and comparable measures, severely affecting social and economic life. Besides improving vaccine designs, booster vaccinations, and the development of adjusted antibodies, it is paramount to identify strategies that might help prevent and treat infections with all current and potential future variants, with a particular urgency for Omicron.

The SARS-CoV-2 Spike protein interacts with very high affinity with cell membrane bound ACE2, followed by a subsequent membrane fusion step. Most neutralizing antibodies from vaccinations and convalescent plasma therapies interfere with the Spike/ACE2 interaction and nearly all therapeutic monoclonal antibodies or nanobodies have been designed to inhibit the binding of Spike to ACE2 [14]. Conceptually, all SARS-CoV-2 variants and “escape mutants” still bind to ACE2 [45-51], which we have recently experimentally shown for the alpha, beta, gamma and delta variants [31]. Moreover, although various other receptors and co-receptors have been proposed, we and others have recently shown that ACE2 is the essential SARS-CoV-2 receptor for respiratory *in vivo* infections using ACE2 mutant mice as well as various human organoids, namely human stem cell derived kidney, gastric, and gut organoids as well as stem cell derived cardiomyocytes [52-54], confirming that ACE2 is of crucial importance for COVID-19 development and the course of the pandemic. ACE2 is a carboxypeptidase which degrades angiotensin II, des-Arg(9)-bradykinin, and apelin, and thereby is a critical regulator of cardiovascular physiology and pathology, kidney disease, diabetes, inflammation, and tissue fibrosis [55]. In addition, the enzymatic activity of ACE2 is protective against acute respiratory distress syndrome (ARDS) caused by viral and non-viral pneumonias, aspiration, or sepsis. Upon infection, both SARS-CoV-2 and SARS-CoV-1 coronaviruses downregulate ACE2 expression, likely associated with the pathogenesis of ARDS[56]. Thus, ACE2 is not only the SARS-CoV-2 receptor but might also play an important role in multiple aspects of COVID-19 pathogenesis and possibly post-COVID-19 syndromes, a hypothesis that has now in part been experimentally confirmed for SARS-CoV-2 induced lung injury using a bacterial orthologue of ACE2 termed B38-CAP [57]. Soluble forms of recombinant human ACE2 are currently utilized by multiple research groups and companies as a potential pan-variant decoy to neutralize SARS-CoV-2 and to supplement the ACE2 carboxypeptidase activity. Our data now show that clinical grade soluble ACE2 can neutralize infections with the SARS-CoV-2 variant Omicron with more than an order of magnitude increased potency when compared to the Wuhan reference strain.

Our modelling data predict that soluble ACE2 could readily associate with the RBD and prefusion trimeric Spike of Omicron, most likely with increased affinity and avidity, which is in line with our experimental data on the markedly improved efficacy of ACE2/APN01 to block Omicron infections. Moreover ACE2/APN01 has very high affinity to all other Variants of Concern and markedly enhanced efficacy to block these infections including the Delta variant, demonstrating that the prediction holds true – clinical grade ACE2 can effectively block all tested SARS-CoV-2 variants and this inhibition is markedly improved against all Variants of Concern and Variants of Interest, including Omicron. APN01 has now undergone phase 2 testing in WHO stage 4-6 COVID-19 patients using intravenous infusions and, in cooperation with researchers at the NIH, we have developed a formulation of APN01 that can be inhaled as an aerosol to directly interfere with the earliest steps of viral infection and COVID-19 development [58]. That this inhalation approach can indeed protect from SARS-CoV-2 infections has been directly confirmed in mice infected with our new mouse adapted SARS-CoV-2 variant, that carries two mutations also found in Omicron [35, 58]. Inhalation of soluble ACE2/APN01 is currently tested in phase 1 trials to assess its safety and tolerability.

The source of the VOC Omicron is currently unclear and multiple hypotheses have been put forward to explain the high number of mutations, including animal hosts as well as protracted infections in immunocompromised hosts that could have led to the gradual evolution of this variant. Additionally, selective pressure by both mass vaccination programs, and antibody and small molecule therapeutics are likely to promote further viral evolution and drive the emergence of therapeutic-/antibody-resistant variant strains of SARS-CoV-2. This viral evolution has already led to the emergence of the Delta and now Omicron VOC that caused devastating global waves of infection and, concerningly, large numbers of re-infections. Our data now shows that clinical grade ACE2/APN01 blocks infectivity of Omicron supporting the notion that this therapeutic is inherently resistant to escape mutations. Our first experimental demonstration that ACE2 inhibits Omicron infections with high efficacy, studies by other groups working on ACE2 and clinical data using soluble ACE2/APN01 support the development of this universal and pan-variant SARS-CoV-2 prevention and therapy. In particular, such an approach should be viable and effective to prevent and treat Omicron infections.

### Limitations of this study

Our study used VeroE6 cells, the classic cellular model for SARS and SARS-CoV-2 infection studies. The study should be expanded to additional cell types as well as human organoids. Moreover, the affinity of Omicron Spike and Omicron RBD should be determined in direct affinity/avidity measurements as well as the impact of non-RBD Spike mutations on the infection process. Of note, from all our previous studies and studies from other groups, the data on soluble ACE2 inhibiting SARS-CoV-2 infections in VeroE6 cells were always supported by results in all other cell types tested. Moreover, we used sera collected 4-6 weeks after the second mRNA vaccination from 4 SARS-CoV-2 naïve healthcare workers, which needs to be expanded to different vaccine regimens and vaccine types and increased sample numbers, though multiple studies are now being released also demonstrating impaired vaccine efficacies to Omicron.

## Acknowledgements

N.M. is supported by the project COV20/00278 from Instituto de Salud Carlos III to N.M. This project has received funding from Ayudas Fundación BBVA a Equipos de Investigación Científica SARS-CoV-2 y COVID-19. J.M.P. has received funding from the T. von Zastrow foundation, the FWF Wittgenstein award (Z 271-B19), the Austrian Academy of Sciences, the Innovative Medicines Initiative 2 Joint Undertaking (JU) under grant agreement No 101005026, and the Canada 150 Research Chairs Program F18-01336 as well as the Canadian Institutes of Health Research COVID-19 grants F20-02343 and F20-02015. A.M. received grant from the Innovative Medicines Initiative 2 Joint Undertaking (JU) under grant agreement No 101005026 and CIMED, Karolinska Institute. J.K has received funding from Knut and Alice Wallenberg foundation/Science for Life Laboratory (SciLifeLab) and Swedish Research Council (2020-05782).

## Materials and Methods

### Viral Strains and isolates

SARS-CoV-2 Wuhan and Omicron strains were isolated from nasopharyngeal swabs from patients in Sweden. The isolates were sequenced by Next-Generation Sequencing (Genbank accession number MT093571).

### Sera of vaccinees

Serum was taken 5-7 weeks after the second immunization with the mRNA vaccine Comirnity (median dose interval 21 days (range 21-24) from four SARS-CoV-2 naïve healthcare workers (75% female, median age 46 [IQR 37-59]) which took part in the COMMUNITY study. The COMMUNITY study was approved by the Swedish Ethical Review Authority (Dnr: 2020-01653).

### Cell lines and cell culture

African green monkey kidney epithelial VeroE6 (ATCC) were grown in Dulbecco’s Modified Eagle’s Medium (DMEM, Thermofisher, supplemented with 1% Non-Essential Amino-Acids (Thermofisher), 10mM HEPES (Thermofisher) and 10% FBS) at 37°C, 5% CO2. Infection and APN01 mediated viral neutralisation assays were conducted at the Karolinska Institute and Karolinska University Hospital.

### Viral neutralization experiments

24h after seeding of VeroE6 cells (5×10^5^ per 48 well), APN01 was mixed with viral particles (MOI of 0.01) of the indicated strains at the given concentrations in DMEM Medium (Thermofischer) containing 5% FBS in 100µl per well and incubated for 30min at 37°C. After the incubation period medium was removed from VeroE6 cells, cells were washed once with PBS to remove any non-attached cells and virus/APN01 mixtures. Cells were incubated with virus for 15h, after which cells were washed 3 times with PBS and lysed with Trizol, subsequently. RNA was extracted using the direct-zol RNA kit (Zymo Research) and assayed by qRT-PCR as previously described (Monteil et al, Cell, 2020) [42]. For serum neutralization assays VeroE6 cells were seeded in 48-well plates as described above 24 hours post-seeding and indicated dilutions of vaccinated subjects sera were mixed with SARS-CoV-2 Wuhan or Omicron strains at an MOI of 0.01 in a final volume of 100ml per well in DMEM (0% FBS) at 37°C under shaking conditions for 30 minutes. The serum dilutions used in this experiment were determined after a neutralization assay against the Wuhan reference strain. After 30 minutes, VeroE6 cells were infected with and Serum/SARS-CoV-2 for 15 hours. 15 hours post-infection, supernatants were removed, cells were washed 3 times with PBS and then lysed using Trizol (Thermofisher) before analysis by qRT-PCR for viral RNA detection as previously described (Monteil et al, Cell, 2020) [42].

### Preparation of recombinant human ACE2

Clinical grade recombinant human ACE2 (amino acids 18-740) was produced by the contract manufacturer Polymun Scientific (Klosterneuburg, Austria) from CHO cells according to GMP guidelines under serum free conditions and formulated as a physiologic aqueous solution, as described previously (Zoufaly et al, 2021, Lancet Respiratory Medicine).

### Visualizations of RBD domains, full-length Spike protein, and Omicron Spike-ACE2 ineractions

Visualizations were rendered with pymol software (the PyMOL Molecular Graphics System, Version 2.0 Schrödinger, LLC), based on a model of the fully glycosylated Spike-ACE2 complex described in Capraz et al. [37] and https://covid.molssi.org//models/#spike-protein-in-complex-with-human-ace2-ace2-spike-binding.

### Primers

The following tables lists the primers used in this study:

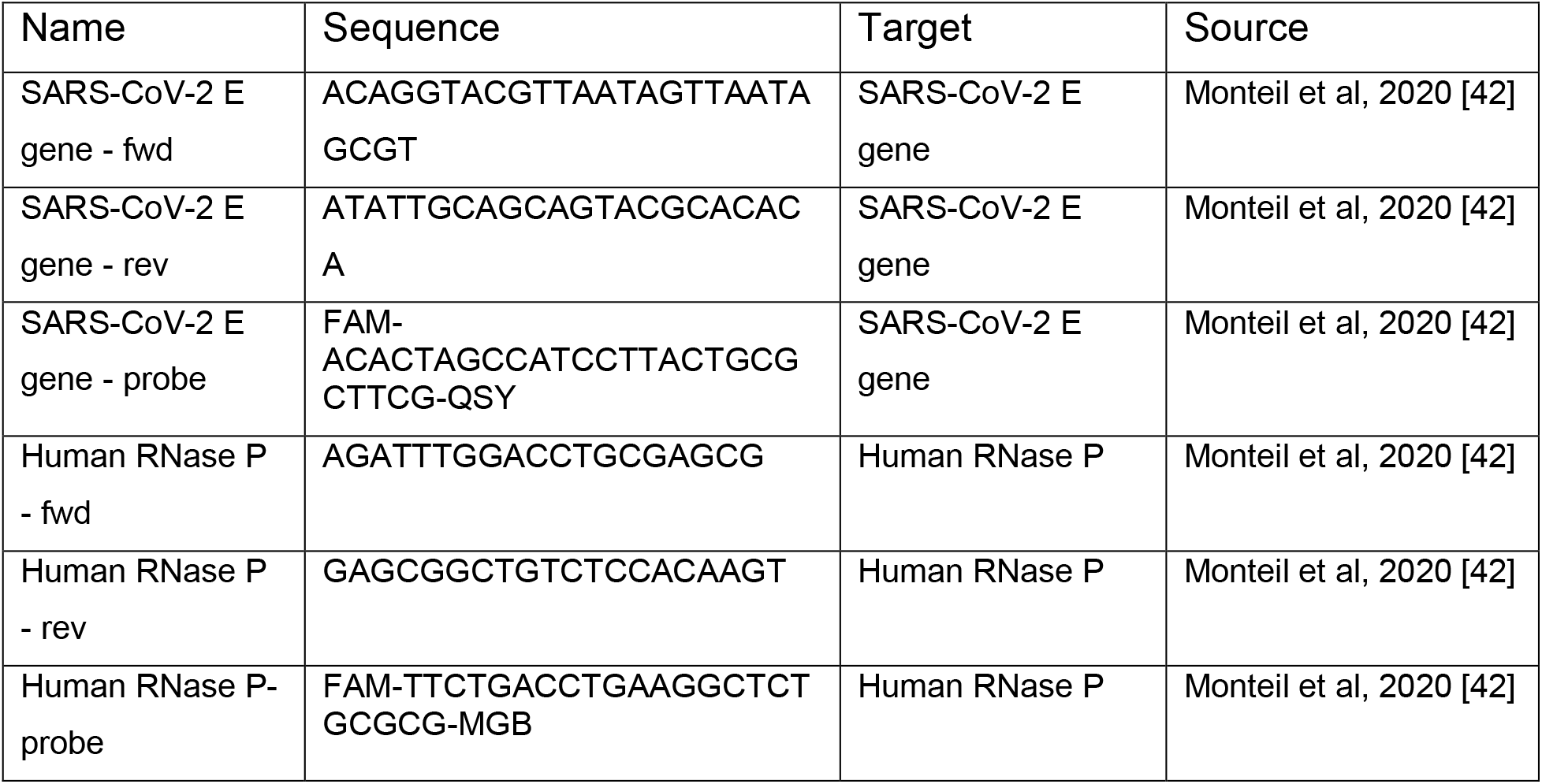

## References

1. Shang, J., et al., Structural basis of receptor recognition by SARS-CoV-2. Nature, 2020. 581(7807): p. 221–224.

2. Zhou, P., et al., A pneumonia outbreak associated with a new coronavirus of probable bat origin. Nature, 2020. 579(7798): p. 270–273.

3. Wang, Q., et al., Structural and Functional Basis of SARS-CoV-2 Entry by Using Human ACE2. Cell, 2020. 181(4): p. 894-904.e9.

4. Hoffmann, M., et al., SARS-CoV-2 Cell Entry Depends on ACE2 and TMPRSS2 and Is Blocked by a Clinically Proven Protease Inhibitor. Cell, 2020. 181(2): p. 271-280.e8.

5. Benton, D.J., et al., Receptor binding and priming of the spike protein of SARS-CoV-2 for membrane fusion. Nature, 2020. 588(7837): p. 327–330.

6. Kyriakidis, N.C., et al., SARS-CoV-2 vaccines strategies: a comprehensive review of phase 3 candidates. npj Vaccines, 2021. 6(1): p. 28.

7. Krammer, F., SARS-CoV-2 vaccines in development. Nature, 2020. 586(7830): p. 516–527.

8. Chen, J., et al., Review of COVID-19 Antibody Therapies. Annual Review of Biophysics, 2021. 50(1): p. 1–30.

9. Svilenov, H.L., et al., Efficient inhibition of SARS-CoV-2 strains by a novel ACE2-IgG4-Fc fusion protein with a stabilized hinge region. bioRxiv, 2020: p. 2020.12.06.413443.

10. Tanaka, S., et al., An ACE2 Triple Decoy that neutralizes SARS-CoV-2 shows enhanced affinity for virus variants. Scientific Reports, 2021. 11(1): p. 12740.

11. Higuchi, Y., et al., Engineered ACE2 receptor therapy overcomes mutational escape of SARS-CoV-2. Nature Communications, 2021. 12(1): p. 3802.

12. Hassler, L., et al., A novel soluble ACE2 protein totally protects from lethal disease caused by SARS-CoV-2 infection. bioRxiv, 2021: p. 2021.03.12.435191.

13. Banerjee, A., K. Mossman, and N. Grandvaux, Molecular Determinants of SARS-CoV-2 Variants. Trends in Microbiology, 2021. 29(10): p. 871–873.

14. Harvey, W.T., et al., SARS-CoV-2 variants, spike mutations and immune escape. Nature Reviews Microbiology, 2021. 19(7): p. 409–424.

15. Planas, D., et al., Reduced sensitivity of SARS-CoV-2 variant Delta to antibody neutralization. Nature, 2021. 596(7871): p. 276–280.

16. Garcia-Beltran, W.F., et al., Multiple SARS-CoV-2 variants escape neutralization by vaccine-induced humoral immunity. Cell, 2021. 184(9): p. 2372-2383.e9.

17. Cele, S., et al., Escape of SARS-CoV-2 501Y.V2 from neutralization by convalescent plasma. Nature, 2021. 593(7857): p. 142–146.

18. Greaney, A.J., et al., Mapping mutations to the SARS-CoV-2 RBD that escape binding by different classes of antibodies. Nature Communications, 2021. 12(1): p. 4196.

19. Lopez Bernal, J., et al., Effectiveness of Covid-19 Vaccines against the B.1.617.2 (Delta) Variant. New England Journal of Medicine, 2021. 385(7): p. 585–594.

20. Jangra, S., et al., SARS-CoV-2 spike E484K mutation reduces antibody neutralisation. The Lancet Microbe, 2021. 2(7): p. e283–e284.

21. Farinholt, T., et al., Transmission event of SARS-CoV-2 Delta variant reveals multiple vaccine breakthrough infections. medRxiv, 2021: p. 2021.06.28.21258780.

22. Christensen, P.A., et al., Delta Variants of SARS-CoV-2 Cause Significantly Increased Vaccine Breakthrough COVID-19 Cases in Houston, Texas. The American Journal of Pathology, 2021.

23. Wolter, N., et al., Early assessment of the clinical severity of the SARS-CoV-2 Omicron variant in South Africa. medRxiv, 2021: p. 2021.12.21.21268116.

24. Bushman, M., et al., Population impact of SARS-CoV-2 variants with enhanced transmissibility and/or partial immune escape. Cell, 2021. 184(26): p. 6229-6242.e18.

25. Eckerle Lance, D., et al., High Fidelity of Murine Hepatitis Virus Replication Is Decreased in nsp14 Exoribonuclease Mutants. Journal of Virology, 2007. 81(22): p. 12135–12144.

26. Eckerle, L.D., et al., Infidelity of SARS-CoV Nsp14-Exonuclease Mutant Virus Replication Is Revealed by Complete Genome Sequencing. PLOS Pathogens, 2010. 6(5): p. e1000896.

27. Kabinger, F., et al., Mechanism of molnupiravir-induced SARS-CoV-2 mutagenesis. Nature Structural & Molecular Biology, 2021. 28(9): p. 740–746.

28. Wilhelm, A., et al., Reduced Neutralization of SARS-CoV-2 Omicron Variant by Vaccine Sera and Monoclonal Antibodies. medRxiv, 2021: p. 2021.12.07.21267432.

29. Schmidt, F., et al., Plasma neutralization properties of the SARS-CoV-2 Omicron variant. medRxiv, 2021: p. 2021.12.12.21267646.

30. Zhang, X., et al., SARS-CoV-2 Omicron strain exhibits potent capabilities for immune evasion and viral entrance. Signal Transduction and Targeted Therapy, 2021. 6(1): p. 430.

31. Wirnsberger, G., et al., Clinical grade ACE2 as a universal agent to block SARS-CoV-2 variants. bioRxiv, 2021: p. 2021.09.10.459744.

32. Gu, H., et al., Adaptation of SARS-CoV-2 in BALB/c mice for testing vaccine efficacy. Science, 2020. 369(6511): p. 1603–1607.

33. Dinnon, K.H., 3rd, et al., A mouse-adapted model of SARS-CoV-2 to test COVID-19 countermeasures. Nature, 2020. 586(7830): p. 560–566.

34. Huang, K., et al., Q493K and Q498H substitutions in Spike promote adaptation of SARS-CoV-2 in mice. EBioMedicine, 2021. 67: p. 103381.

35. Gawish, R., et al., ACE2 is the critical &lt;em&gt;in vivo&lt;/em&gt; receptor for SARS-CoV-2 in a novel COVID-19 mouse model with TNF-and IFNγ-driven immunopathology. bioRxiv, 2021: p. 2021.08.09.455606.

36. Hoffmann, D., et al., Identification of lectin receptors for conserved SARS-CoV-2 glycosylation sites. The EMBO Journal, 2021. 40(19): p. e108375.

37. Capraz, T., et al., Structure-guided glyco-engineering of ACE2 for improved potency as soluble SARS-CoV-2 decoy receptor. bioRxiv, 2021: p. 2021.08.31.458325.

38. Aggarwal, A., et al., SARS-CoV-2 Omicron: evasion of potent humoral responses and resistance to clinical immunotherapeutics relative to viral variants of concern. medRxiv, 2021: p. 2021.12.14.21267772.

39. Cameroni, E., et al., Broadly neutralizing antibodies overcome SARS-CoV-2 Omicron antigenic shift. bioRxiv, 2021: p. 2021.12.12.472269.

40. Cao, Y.R., et al., Omicron escapes the majority of existing SARS-CoV-2 neutralizing antibodies. bioRxiv, 2021: p. 2021.12.07.470392.

41. Planas, D., et al., Considerable escape of SARS-CoV-2 variant Omicron to antibody neutralization. bioRxiv, 2021: p. 2021.12.14.472630.

42. Monteil, V., et al., Inhibition of SARS-CoV-2 Infections in Engineered Human Tissues Using Clinical-Grade Soluble Human ACE2. Cell, 2020. 181(4): p. 905-913.e7.

43. Monteil, V., et al., Human soluble ACE2 improves the effect of remdesivir in SARS-CoV-2 infection. EMBO Molecular Medicine, 2021. 13(1): p. e13426.

44. Karim, S.S.A. and Q.A. Karim, Omicron SARS-CoV-2 variant: a new chapter in the COVID-19 pandemic. The Lancet, 2021. 398(10317): p. 2126–2128.

45. Gobeil Sophie, M.C., et al., Effect of natural mutations of SARS-CoV-2 on spike structure, conformation, and antigenicity. Science. 373(6555): p. eabi6226.

46. Tchesnokova, V., et al., Acquisition of the L452R Mutation in the ACE2-Binding Interface of Spike Protein Triggers Recent Massive Expansion of SARS-CoV-2 Variants. Journal of Clinical Microbiology. 59(11): p. e00921–21.

47. Tian, F., et al., N501Y mutation of spike protein in SARS-CoV-2 strengthens its binding to receptor ACE2. eLife, 2021. 10: p. e69091.

48. Zhou, D., et al., Evidence of escape of SARS-CoV-2 variant B.1.351 from natural and vaccine-induced sera. Cell, 2021. 184(9): p. 2348-2361.e6.

49. Ou, J., et al., V367F Mutation in SARS-CoV-2 Spike RBD Emerging during the Early Transmission Phase Enhances Viral Infectivity through Increased Human ACE2 Receptor Binding Affinity. Journal of Virology. 95(16): p. e00617–21.

50. Motozono, C., et al., SARS-CoV-2 spike L452R variant evades cellular immunity and increases infectivity. Cell Host & Microbe, 2021. 29(7): p. 1124-1136.e11.

51. Cai, Y., et al., Structural basis for enhanced infectivity and immune evasion of SARS-CoV-2 variants. Science, 2021. 373(6555): p. 642–648.

52. Garreta, E., et al., A diabetic &lt;em&gt;milieu&lt;/em&gt; increases cellular susceptibility to SARS-CoV-2 infections in engineered human kidney organoids and diabetic patients. bioRxiv, 2021: p. 2021.08.13.456228.

53. Beumer, J., et al., A CRISPR/Cas9 genetically engineered organoid biobank reveals essential host factors for coronaviruses. Nature Communications, 2021. 12(1): p. 5498.

54. Iwanski, J., et al., Antihypertensive drug treatment and susceptibility to SARS-CoV-2 infection in human PSC-derived cardiomyocytes and primary endothelial cells. Stem Cell Reports, 2021. 16(10): p. 2459–2472.

55. Kuba, K., et al., Trilogy of ACE2: A peptidase in the renin–angiotensin system, a SARS receptor, and a partner for amino acid transporters. Pharmacology & Therapeutics, 2010. 128(1): p. 119–128.

56. Kuba, K., T. Yamaguchi, and J.M. Penninger, Angiotensin-Converting Enzyme 2 (ACE2) in the Pathogenesis of ARDS in COVID-19. Frontiers in Immunology, 2021. 12(5468).

57. Yamaguchi, T., et al., ACE2-like carboxypeptidase B38-CAP protects from SARS-CoV-2-induced lung injury. Nature Communications, 2021. 12(1): p. 6791.

58. Shoemaker, R.H., et al., Development of a novel, pan-variant aerosol intervention for COVID-19. bioRxiv, 2021: p. 2021.09.14.459961.

